# New insights into the paradoxical distribution of *IRC7* in *Saccharomyces cerevisiae* and its associated phenotypic and genomic landscapes

**DOI:** 10.1101/2020.09.24.307892

**Authors:** Javier Ruiz, Miguel de Celis, María Martín-Santamaría, Iván Benito-Vázquez, Ana Pontes, Val F. Lanza, José Paulo Sampaio, Antonio Santos, Ignacio Belda

**Affiliations:** Department of Genetics, Physiology and Microbiology. Unit of Microbiology. Biology Faculty, Complutense University of Madrid, 28040, Madrid, Spain; Departamento de Ciências da Vida, Faculdade de Ciências e Tecnologia, Universidade Nova de Lisboa, 2829-516 Caparica, Portugal; Department of Microbiology, Ramón y Cajal University Hospital, IRYCIS, 28034, Madrid, Spain

**Author notes:** Corresponding authors: **Antonio Santos**, Email address, **Ignacio Belda**, Email address.

**Keywords:** Saccharomyces cerevisiae, IRC7, Domestication, Phenotyping, Genomic survey, Wine clade

## Abstract

Biotic and abiotic factors of wine fermentations have led to the accumulation of numerous genomic hallmarks of domestication in *Saccharomyces cerevisiae* wine strains. Here we have studied the paradoxical distribution of a dominant allele of *IRC7* in wine yeast strains. This gene encodes a cysteine-S-β-lyase and presents two alleles: a wild full-length allele (*IRC7^F^*) and a mutated one (*IRC7^S^*), harboring a 38bp-deletion. Interestingly, *IRC7^S^*-coding for a less active enzyme-appears in the great majority of wine strains. Studying its global distribution among phylogenetic clades, we observed that *IRC7^S^* allele is dominant just in wine strains, having a moderate presence in other domesticated clades (Beer, Bread and Wine-PDM), but being completely absent in wild clades, appearing as a new hallmark of domestication. To explain this paradoxical distribution, we performed an *IRC7*-rooted phenotypic-wide survey, demonstrating that *IRC7^S^*-homozygous (HS) wine strains have both fitness (lower lag phases and higher growth rates) and competitive (killer toxin resistance, pseudohyphal growth) advantages. Hence, we performed a genome-wide survey across domesticated clades, finding a set of mutations that are conserved among wine strains and potentially associated to *IRC7^S^* allele, which can help to explain the outstanding phenotype of HS strains and their dominant distribution in wines.

**Originality-Significance Statement:** *S. cerevisiae* is one of the best studied microbes due to its industrial importance and its use as a eukaryotic model organism. *S. cerevisiae* is also an interesting model for studying the effects of domestication in yeast genomic, phenotype and ecology. The study of how wild *S. cerevisiae* strains evolved into a greatly adapted domesticated strains and changed its lifestyle drastically is still of great interest. In the case of wine populations, strains have accumulated numerous hallmarks of domestication in their genome, related with their great phenotypic adaptation to this environment. Here, we report a new hallmark of domestication in wine strains; the *IRC7* deleted allele (*IRC7^S^*) and present the first insights about its unexpected global distribution among phylogenetic clades, understanding the genomic context and the phenotypic implications of this allele in wine strains.

## Introduction

*Saccharomyces cerevisiae* is a eukaryotic model microorganism used in cellular physiology, molecular biology and genetics. However, much is still unknown about its metabolism in natural environments, beyond the laboratory (Liti *et al.*, 2015). *S. cerevisiae* strains are worldwide distributed, occurring in multiple wild substrates and associated to several human activities (Wang *et al.*, 2012). From its natural origin, the genome and the phenome of *S. cerevisiae* strains have been shaped for millennia, through different domestication events driven by a combination of natural and anthropic selection forces. This has originated highly-adapted strains, defining different -wild and domesticated-lineages of *S. cerevisiae* during its evolutionary history (*domesticated clades:* Wine, Wine PDM (Flor), Beer1, Beer2, Bread and Sake; *wild clades:* West African, Mediterranean Oak, North America & Japan and Malaysia) (Gallone, *et al.*, 2016; Gonçalves *et al.*, 2016; Coi *et al.*, 2017). These clades reflect, not only their geographical distribution, but also their lifestyle in association to different niches (Liti *et al.*, 2009; Schacherer *et al.*, 2009; Strope *et al.*, 2015, Peter *et al.*, 2018). Because of the selective pressures of the different niches and, as a consequence of their high genome plasticity, *S. cerevisiae* strains are highly adapted to each environment they inhabit (Legras *et al.*, 2018).

One of these well-defined monophyletic lineages is the Wine clade, including isolates from wine must, grapes and vineyard soils (Almeida *et al.*, 2017). It has been demonstrated that this clade arose from a single bottleneck event of domestication (Peter *et al.*, 2018), being Mediterranean Oak population the proposed wild origin for wine yeasts clade (Almeida *et al.*, 2015). After that, viticulture expansion through Europe and Mediterranean Sea led to the migration of yeasts associated to grapevines to all the wine-producing regions around the world (Legras *et al.*, 2007). Since, probably, more than 9.000 years (McGovern *et al.*, 2004), the challenging conditions of wine fermentation and the human pressure to achieve wine desirable traits, have forced wine strains to a great adaptation, implying important changes on its genome if compared to non-wine strains (Marsit & Dequin, 2015). Therefore, numerous hallmarks of domestication have been reported in wine strains, as examples of the adaptive process to wine environment (Belda *et al.*, 2020).

In this work, we report new genetic and phenotypic signatures within the wine yeasts population. These potential hallmarks of domestication appeared associated to the previously reported paradoxical distribution of *IRC7* alleles in wine yeasts (Belda *et al.*, 2016). This gene, coding for a cysteine-S-conjugate β-lyase (EC 4.4.1.13), is the direct responsible for the release of volatile thiols (3SH (3-sulfanylhexan-1-ol) and 4MSP (4-methyl-4-sulfanylpentan-2-one)) from their cysteinylated precursors, releasing pleasant aromas in white wines (Harsch & Gardner, 2013; Howell *et al.*, 2005; Swiegers & Pretorius, 2007; Tominaga *et al.*, 1998). Two alleles have been reported for *IRC7* in *S. cerevisiae*: a 1200-bp wild type full-length allele (*IRC7^F^*) encoding for a 400 amino acid protein, and an altered allele carrying a 38-bp deletion (*IRC7^S^*), appearing a premature stop codon, and thus encoding for a shorter (360 amino acids) and less functional enzyme (Roncoroni *et al.*, 2011). Hence, three *IRC7* genotypes have been described in *S. cerevisiae* strains: homozygous strains for the full-length *IRC7* allele (HF), heterozygous strains (HT), and homozygous strains for the short-length *IRC7* allele (HS). Surprisingly, the great majority of *S. cerevisiae* wine strains harbor -in homozygosis-the *IRC7^s^* allele, and therefore, they have a less functional β-lyase enzyme to release aromatic thiols (Roncoroni *et al.*, 2011; Belda *et al.*, 2016; Cordente *et al.*, 2019).

To explain the remarkable high presence of HS-strains in wine environment, we propose an extensive genomic and phenomic study to seek for other metabolic and genetic traits associated to *IRC7* genotypes. Thus, we performed (1) an *IRC7* genotyping survey in a global collection of *S. cerevisiae* strains, isolated from different origins and niches. With the aim of understand the global distribution of *IRC7* alleles in different phylogenetic clades we have conducted (2) a high throughput phenotyping study to seek for metabolic and growth differences between the three *IRC7* genotypes, and (3) a genome wide association study of *IRC7*-related genetic variants, looking for complementary genomic patterns that match, and help to understand, the singular *IRC7* allele distribution among wine strains.

## Results & Discussion

### IRC7 Allele Distribution in S. cerevisiae Populations

Evolutionary history of *S. cerevisiae* has drawn a phylogenetic tree where domesticated and wild clades cluster into two well-defined groups, with the exceptions of Mediterranean Oak and Sake clades (Peter *et al.*, 2018). Meanwhile wild clades represent different geographical origins (Mediterranean Oak, North America & Japan, West Africa, Philippines and Malaysia), domesticated clades grouped based on their anthropological niche (wine, beer, bread and sake). It can be observed an evolutionary progress from Asiatic wild populations to domesticated yeast population. These domestication events are accompanied with specific phenotypic traits that come from genetic variants, from Single-Nucleotide Polymorphism to Copy Number Variation or Horizontal Gene Transfer (Belda *et al.*, 2020).

Figure 1 shows the distribution of *IRC7* alleles among the different *S. cerevisiae* phylogenetic clades. For this gene, the two alleles described by Roncoroni *et al.* (2011) (*IRC7^F^* and *IRC7^S^*) defined three genotype groups: HF for homozygous strains for *IRC7^F^* allele; HT for heterozygous strains; and HS for homozygous strains for *IRC7^S^* allele. To study how the *IRC7* alleles are distributed in *S. cerevisiae* populations, an *IRC7* genotyping study using a collection of 283 *S. cerevisiae* strains, representing different phylogenetic clades and origins (Supplementary File S1) was performed.

**Figure 1:**
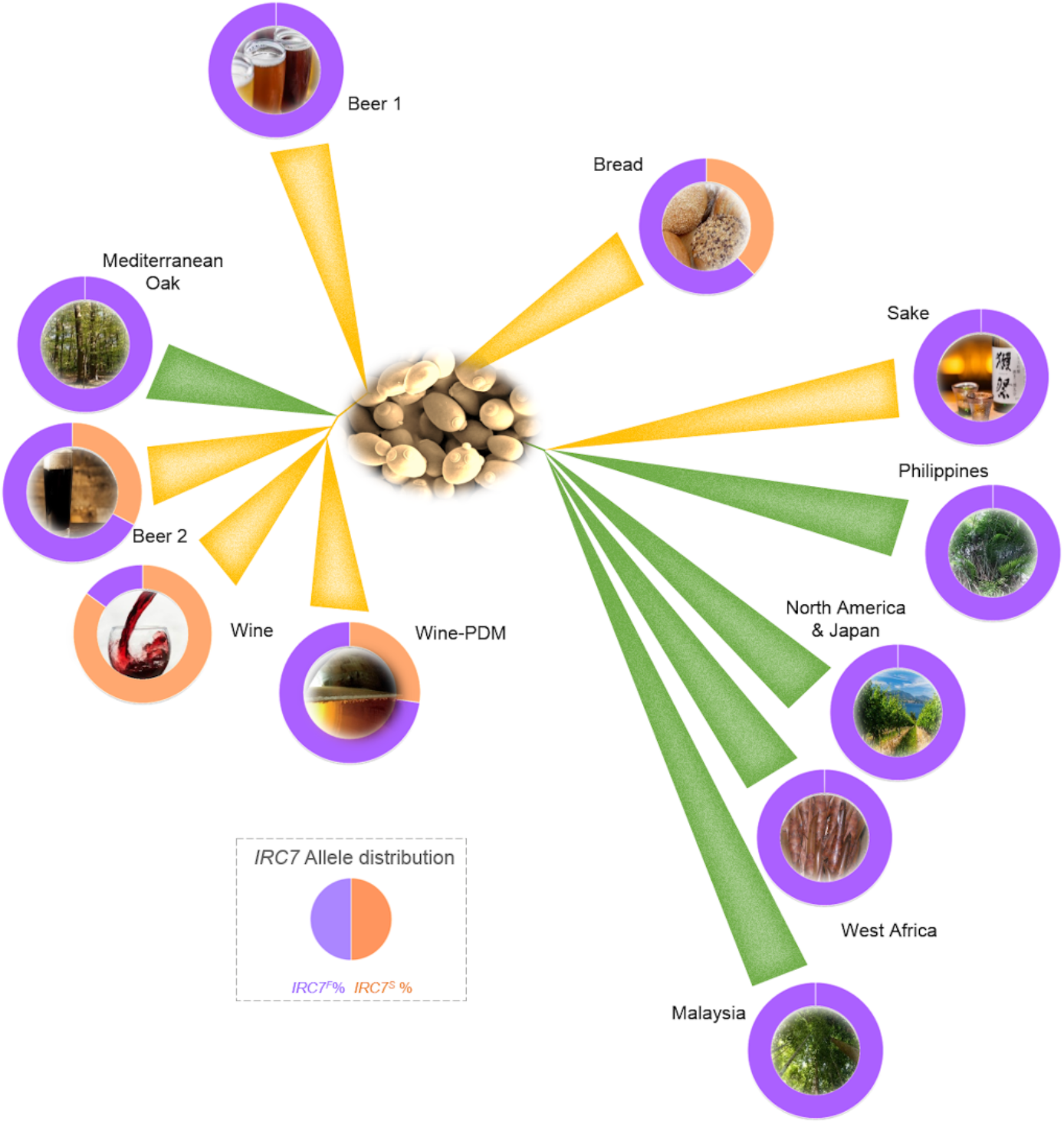
Phylogenetic scheme of *S. cerevisiae* populations including the *IRC7* allele distribution (orange for *IRC7^S^* and purple for *IRC7^F^*) of each phylogenetic clade, using a dataset of 283 strains (Supplementary File S1). Yellow branches highlight domesticated clades and green branches highlight wild clades.

As reported before (Roncoroni *et al.*, 2011, Belda *et al.*, 2016, Cordente *et al.*, 2019), we observed that the *IRC7^S^* was the dominant allele in the wine clade (85%). Here we conclude that there was no regional prevalence in the distribution of *IRC7* alleles, as, although it was possible to isolate strains harboring *IRC7^F^* allele in different wine regions, *IRC7^S^* allele was dominant among wine strains independently on their origin (Supplementary File S1). However, here we found that *IRC7^S^* allele was also present in a few strains from other domesticated clades, but it was completely absent in the wild clades. Domesticated clades such as Wine-PDM (*“Prise de Mousse”,* a strain family mainly represented by Flor wine strains (Legras *et al.*, 2014)) and Beer2, both defined as intermediate groups between non-wine and wine strains (Dunn *et al.*, 2012; Gallone *et al.*, 2016), showed an intermediate presence of the *IRC7^S^* allele (27.3% and 35.0%, respectively). The intermediate presence of both *IRC7* alleles was also observed in the domesticated clade of Bread strains, with a 37.5% of the strains harboring the *IRC7^S^* allele. All the strains pertaining to the other domesticated (Beer 1 and Sake), and wild clades (Mediterranean Oak, Philippines, North America & Japan, West Africa and Malaysia) only harbored the *IRC7^F^* wild-type allele.

These results suggest that *IRC7^S^* allele might represents a new hallmark of domestication in certain domesticated clades of *S. cerevisiae*, with an outstanding presence in wine strains. This hypothesis is supported by the fact that other species, out of the *Saccharomyces* genus, as *Torulaspora delbrueckii,* do not harbor the orthologous *IRC7^S^* allele in any of the strain checked (Belda *et al.*, 2017). This result would contravene the intuition derived from artificial selection processes, as *IRC7^F^* is strongly related with the production of pleasant aromas in wine fermentations, but the strains harboring the *IRC7^S^* allele (inefficient for aromatic thiols releasing) are dominant in the wine clade. Thus, we aimed to understand the biological basis of this paradoxical allele distribution, still a pending task for wine yeast researchers (Belda *et al.*, 2016, Santiago & Gardner, 2015, Cordente *et al.*, 2019).

### Phenotypic landscape based on IRC7-genotype

As aforementioned, a truncated and less-functional version of the β-lyase enzyme encoded by *IRC7^s^* is widespread among wine yeast strains. To understand the biological basis of its selection as a hallmark of domestication here we carried out a high throughput phenotyping study looking for growth differences between the defined *IRC7-*genotype groups, beyond its cysteine-S-conjugate β-lyase activity. Thirty *S. cerevisiae* strains, representing the 3 different *IRC7* genotypes (10 HF, 10 HT, and 10 HS) (Table S1) were assayed in a panel of 48 culture media and conditions (testing different carbon and nitrogen sources, physic-chemical conditions and antimicrobials) (Table S2). Growth curves in axenic cultures were analysed to obtain growth parameters (lag phase, growth rate and efficiency) (Supplementary File S2).

Figure 2A represents normalized average values of the growth parameters obtained in all the growth conditions tested for the group of strains pertaining to each *IRC7* genotype. We observed a high diversity between *S. cerevisiae* strains in lag phase length, as reported by Ferreira *et al.* (2017). Likewise, growth rate was also very variable among the strains. Despite this variability, HF-strains showed a generalized lower fitness, exhibiting longer lag phases and lower growth rates in most of the growth conditions tested. Efficiency was similar among the *IRC7* genotypic groups. Thus, we observed clustered growth patterns highly associated to *IRC7-genotype* groups (Figure S1), where HS-strains exhibited an increased overall fitness that may imply a competitive advantage for dominating the microbial succession established during wine fermentations, as we explored later on.

**Figure 2:**
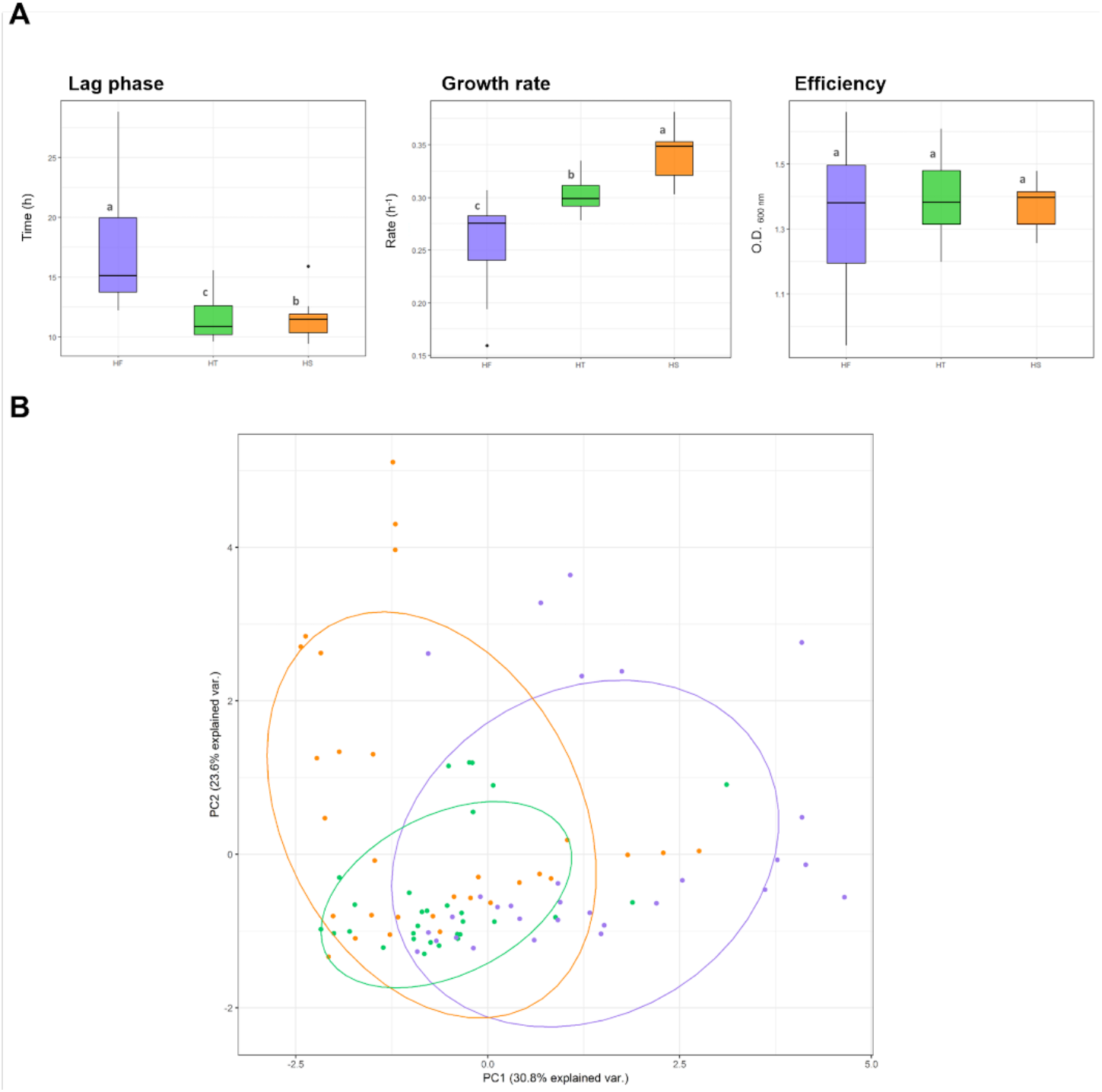
**A** Boxplot of the average lag time, growth rate and efficiency of the three different strains groups, according to their *IRC7* genotype. The performance in all the media panel were considered together in each plot. Different letters indicate the existence of statistical differences (p < 0.05). **B** Principal Component Analysis (PCA) representation of the wine fermentative parameters measured after the fermentation of the strains in synthetic grape must. Principal Component 1 represents the 30.8% of the variance and presents statistically differences among the HS and HF groups (P<0.01).

In addition, metabolite profiles in wine fermentations can display other subtle differences between strains with different genotypes (Legras *et al.*, 2018). Thus, we performed microvinifications in synthetic grape must to evaluate strain-associated patterns in basic wine parameters at the end of the alcoholic fermentation. Figure 2B shows an ordination analysis of the global metabolite patterns obtained in wine fermentations performed with all the strains tested, highlighting the *IRC7* genotype-associated clusters. PC1 represents the 31% of the variance and shows statistically differences between HS and HF groups (p<0.01). Thus, in addition to the different fitness patterns observed in growth parameters (Figure 2A), yeast strains produced different metabolic profiles in wine fermentations, with a certain dependence on the *IRC7* genotype. Considered together, these data suggest that the phenotypic behavior of *S. cerevisiae* strains could be associated to the *IRC7* genotype.

### Competitive ability of strains based on IRC7-genotype

When inoculated on grape must, yeast cells have to adapt to a highly stressful and competitive environment, performing a switch between respiration and fermentation metabolism, which is the main factor determining lag phase duration (Vermeesch *et al.*, 2019). Therefore, the quick adaptation to these conditions that leads to the beginning of the exponential growth phase will be decisive for the subsequent population to compete efficiently with the microbial populations that inhabit the same niche. Thus, longer lag phases and lower growth rates can strongly determine the worse implantation of the strains on fermentative environments, even though the strains have the same biomass production yield, i.e. efficiency.

To evaluate the potential advantage of HS-strains to colonize wine fermentations as a consequence of a fitness-related increased competitive ability, we carried out yeast-to-yeast (HF vs HS) competition experiments under fermentative and non-fermentative conditions. We selected 3 strains representing HS genotypes and 4 representing HF genotypes. In competition assays, each HS strain was co-inoculated, at the same cellular concentration, with one of each HF-strain. Figure 3A shows the better competitive profiles obtained for HS-strains, independently on the growth environment, with averaged implantation percentages of 65%-82%-75% and success rates of 67%-100%-83% in wine, beer and non-fermentative conditions, respectively. As we showed before, HF-strains exhibited lower fitness in most of the tested media and conditions (Figure 2A). This disadvantage, caused by longer lag phases and lower growth rates, is one of the major determinants of competitive fitness in multi-strains environments (Schmidt *et al.*, 2019), and can explain the difficulty of HF-strains to dominate in mixed populations at the end of fermentation assays (Figure 3A).

**Figure 3:**
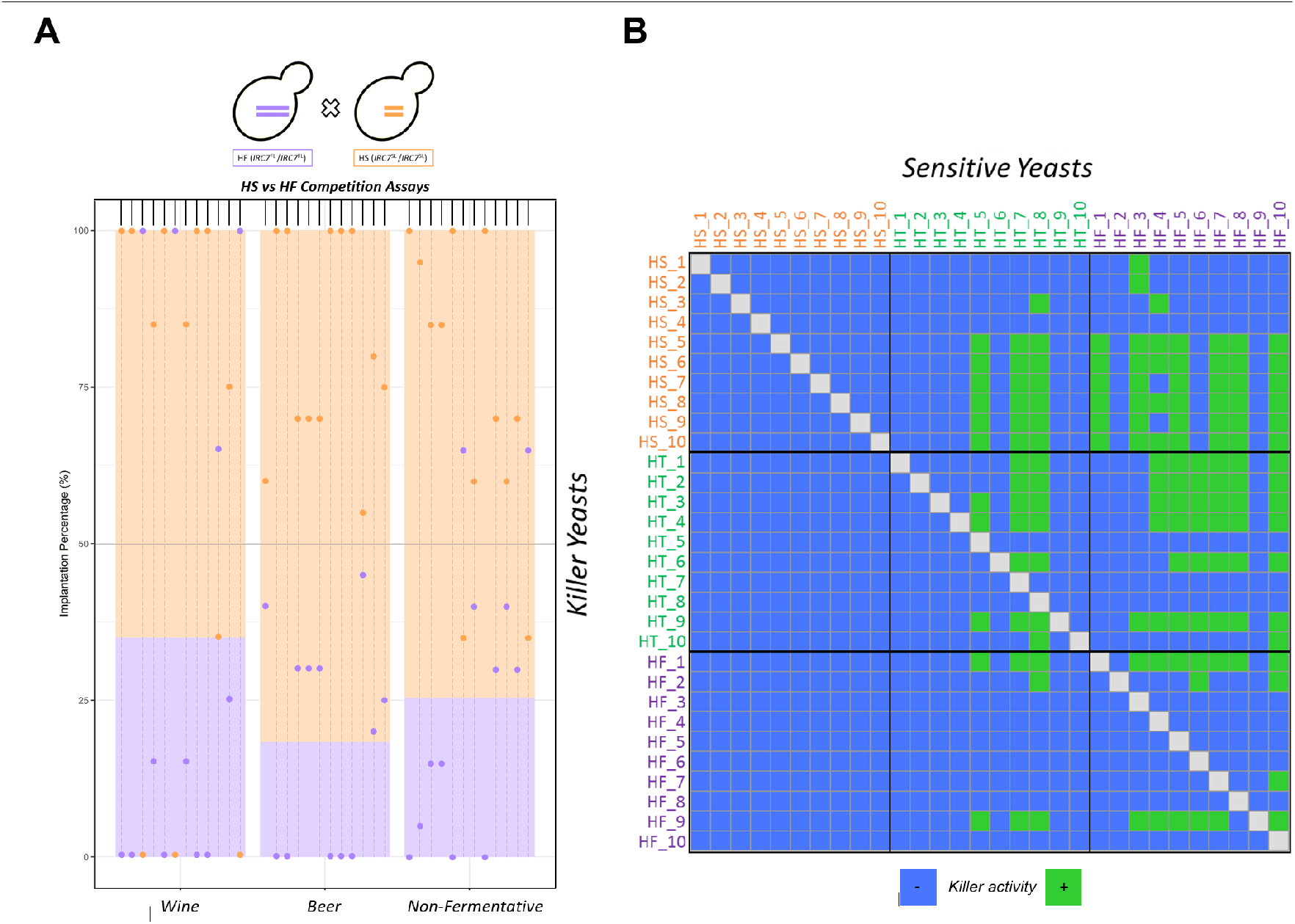
(**A)** Competition assays of HS and HF-strains in wine, beer and non-fermentative mimic media. Twelve combinations of HS and HF-strains (1:1 proportion) were assayed in the three different conditions. Implantation percentage after seven days of culture, analyzed by interdelta patterns typing, is representing. Bar plots indicate the average implantation of each *IRC7* group and dot plots indicate the implantation percentage of each competition assay. (**B)** Killer activity of the yeast crosses assays of thirty strains representing each IRC7-genotype group (HS, orange, HT, green and HF purple) is represented. Killer yeasts term refers to the yeast tested for killer activity, and sensitive yeasts refers to yeast tested for killer sensibility. Yellow color indicates that we observed killer activity in that cross assay. Blue color indicates that we did not observe killer activity.

In addition to the lower fitness of HF-strains at the beginning of the fermentation, there are other facts that may determine the implantation of the strains in competitive environments such as wine fermentations. Some *S. cerevisiae* strains are able to secret killer toxins, active against other sensitive yeasts including strains of the same species. Killer strains can control the implantation of other strains by producing killer toxins, as a great advantage in competitive environments as wine fermentation (Marquina *et al.*, 2002). We evaluated the killer phenotype in our studied strains collection, looking for patterns in killer activity and/or resistance associated with the *IRC7* genotype. Figure 3B shows the results obtained for the killer crosses assays, representing the sensitivity/resistance phenotypes of the strains tested. Killer phenotype seems to be related with the *IRC7-genotype* of *S. cerevisiae* strains, as all HS-strains tested were resistant to the strains tested as potential killer strains. Contrary, 7 of 10 HT-strains and only 2 of 10 HF-strains were killer resistant against the strains tested. In addition, killer activity (against at least one of the sensitivity strains tested) was observed in 9 of 10 HS-strains, 7 of 10 HT-strains, and only in 4 of 10 HF strain. As far as our knowledge reaches, Irc7p enzyme is not directly nor indirectly involved in killer toxins production neither killer resistance, we have clear indicatives that other metabolic activities, coming from other genetic variants, may be associated with the *IRC7* genotype.

All these phenotypic results demonstrated that *IRC7-*genotype can be associated, in some way, to the phenotype behavior of *S. cerevisiae* strains, showing an overall advantage of HS-strains in most of the experiments performed. Thus, these results may explain the great dominance of HS-strains in wine clades, but the reason why HS strains were not strictly dominant in other highly domesticated clades like Flor (Wine-PDM), Beer or Bread, remains to be answered (Figure 1). One explanation is that these clades, in comparison to Wine clade, are inhabiting well controlled fermentation processes where the strains used are not exposed to a highly competitive pressure and without the presence of other yeast strains or microbial species, as it occurs in grape musts (Conacher *et al.*, 2019). Secondary wine, beer and bread fermentations generally represent low diverse environments, therefore, the competitive advantage that we have demonstrated in HS-strains, may not be an important fact compared to wine environment. These results led us to look for specific mutations or sequence variants that were conserved among wine HS-strains, that can help us in explaining the advantageous competitive phenotype of these strains in such an environment.

### IRC7-rooted genomic survey

#### Identification of genetic variants underlying the HS-strains associated phenotype

The above-mentioned results indicated that *IRC7* allele in *S. cerevisiae* strains is somehow associated with overall aspects of yeast phenotype, in terms of growth performance and yeast-yeast competition ability. In order to explain the genomic bases of this outstanding phenotype of HS-strains, we carried out a whole-genome sequencing of 9 of our studied strains, including 3 representative strains (HS4, HS6, HS9; HT3, HT6, HT10; HF1, HF2, HF9) of each *IRC7-genotype* and a further variant calling analysis to identify gene variants potentially associated to *IRC7^S^* genotype. We used *S. cerevisiae* VL3 strain as the reference genome for variant calling, as it is a well-studied wine strain homozygous for the wild-type *IRC7^F^* gene. The presence and co-occurrence of mutations among the studied strains were represented in a bipartite network, including moderated and highly important mutations, and discarding those widespread mutations found in all the strains and those rare ones just found in one single strain (Figure 4). To focus our analysis on the potential genomic patterns associated to *IRC7* alleles, we studied in detail those mutations conserved among all the HS-strains and absent in all HF-strains (highlighted in red in Figure 4A). With this premise, HS-strains shared 39 mutations affecting to 13 different genes (Table S3, Figure 4B). These mutations, as were completely absent in the studied HF-strains, were considered potential candidates for helping us to understand its marked clade-associated distribution and the fitness advantages found in the studied *IRC7*-HS wine strains.

**Figure 4:**
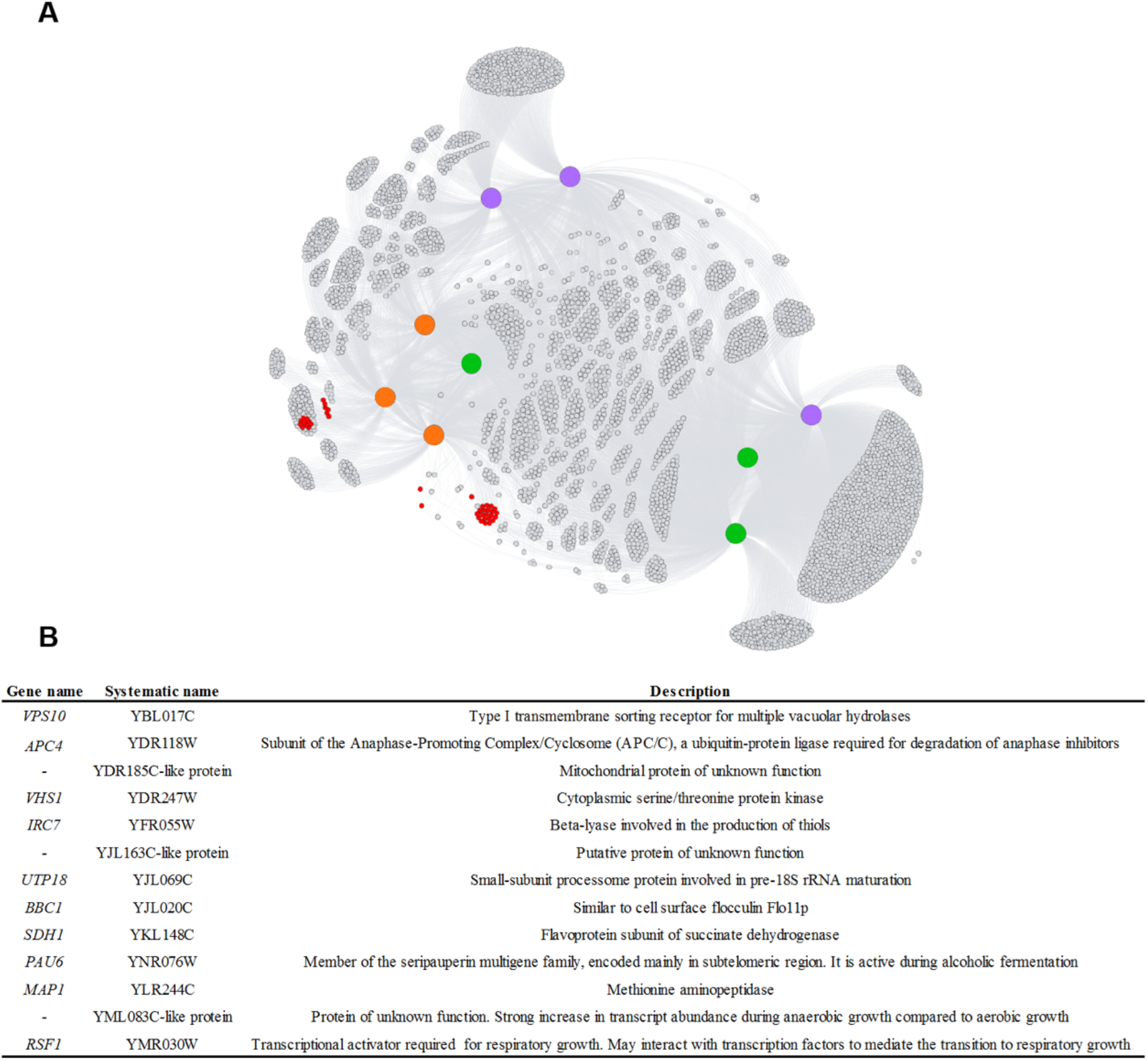
**A** Bipartite network representation of high important mutations detected in a variant calling analysis using *S. cerevisiae* VL3 genome as reference. Mutations present in all the strains and those present only in one strain were discarded. Those mutations present in all the 3 HS strains but absent in HF strains are highlighted as red nodes. Orange hubs represent HS strains, green hubs represent HT strains, and purple hubs represents HF strains. **B** Genes containing mutations found in the three HS-strains and absent in the three HF strains (red dots in **A**).

Among the identified mutations, we obviously found the above-described *IRC7* 38-bp deletion. Irc7p may have an implication in *S. cerevisiae* growth under some specific conditions, due to their role in cysteine and methionine metabolisms. However, the dominant presence of *IRC7*-HS strains in Wine clade (Figure 1) and their notable growth and competitive advantages reported in this work (Figure 2) suggest that *IRC7* might have some additional metabolic traits with a higher implication in the overall fitness. Santiago & Gardner (2015) proposed a putative role of *IRC7^F^* coding enzyme on cysteine homeostasis by demonstrating its cysteine desulfhydrase activity. A fully functional Irc7p (encoded by the *IRC7^F^* allele) could compromise the availability of the intracellular cysteine pool, therefore, due to the important impact of cysteine on glutathione production, *IRC7* could have a relevant role in the protection against oxidative damage in fermentation environments (García-Ríos & Guillamón, 2019).

Based on the foregoing, we performed an oxidative stress shock assay to test the response of our studied strains. In the conditions tested, and considering the average oxidative stress level of the different *IRC7-*genotype groups, HS-strains showed a slight lower oxidative damage levels if compared to HF-strains (Figure S2A). In addition, *IRC7* is regulated by copper availability (*IRC7* expression is inhibited under high copper conditions) and it may have a role on copper tolerance as it has been reported (Cordente *et al.*, 2019). Copper compounds are commonly used in vineyards as phytosanitary agents to avoid negative sulfur aromas during winemaking (Swiegers & Pretorious, 2007). Cordente *et al.*, (2019) proposed that an Irc7p enzyme with a reduced ability to degrade cysteine ensure a higher intracellular cysteine pool, needed for the synthesis of cysteine-rich copper metallothioneins, as Cup1p. Thus, the effect of copper on the growth ability of our strains was tested. Figure S2B represents the growth parameters of each *IRC7*-genotype group in the SD medium supplemented with 375 μM of CuCl2. HS-strains showed an overall better growth performance in the copper supplemented medium, exhibiting, on average, shorter lag phases and higher growth rates if compared to HF-strains.

Despite these outcomes, other genetic variants, non-related with *IRC7* but associated with HS-strains, may help to explain the definitory phenotype that we found for HS-strains. Among these mutations, found in our HS-strains studied in detail at the genomic level, we stand out those affecting *VPS10, PAU6* and *RSF1* genes. *VPS10* contains 24 mutations in the three HS-strains that are not found in any of the three HF-strains studied, 5 of them even lacked in the three HT-strains. Vps10p is a type I membrane receptor, involved on trafficking of carboxypeptidase Y (CPY) and other vacuolar hydrolases to the vacuole (Bowers & Stevens, 2005). Serviene *et al.*, (2012) demonstrated that the deletion of *VPS10* led to a K2 killer toxin hypersensitivity caused by a defect in cell osmoregulation. This role of Vps10p on killer resistance lead us to hypothesize the potential impact of the *VPS10* version found in *IRC7-*HS-strains in determining their strong and widespread resistance to killer toxins activity (Figure 3B). *PAU6* is a member of the large PAU family, which might have a specific role in the adaptation of yeasts to environmental stress and in the fitness of yeast during alcoholic fermentation (Luo & van Vuuren, 2009), and in the case of *PAU5,* a direct role in the resistance against yeast killer toxins has been observed (Rivero *et al.*, 2015). The biological role of *PAU6* is still unknown, but the high percentage identity (84%) between Pau6p and Pau5p allows us to hypothesize that may also be involved in killer toxins resistance.

*RSF1* encodes for a transcriptional factor required for the transition to respiratory growth. It is specifically necessary for the use of glycerol and ethanol as carbon sources (Liu *et al.*, 2005), and it is also involved in the sporulation process. In fact, it has been reported that the polymorphism we identified here (XIII_93636_C, D181G), reduces the function of this gene, and therefore, the efficiency of sporulation (Gerke *et al.*, 2009). As sporulation and filamentous growth are conflicting behaviors (Cullen *et al.*, 2012), and the latter gives cells an advantage in food foraging conditions, we checked the pseudohyphal growth ability of our strain panel in SLAD medium (stablishing nitrogen nutrient limitations). The pseudohyphal growth was observed in all the HS-strains, but only in 30% of HF-strains (Figure S3). This supports the potential association between the genetic variant of *RSF1* gene and the pseudohyphal growth ability of yeasts, mainly shown by HS-strains, which may have an implication in their dominance and improved fitness in wine fermentations.

This genomic comparison allowed us to identify mutations associated to *IRC7^s^* allele, helping us to understand the genomic determinants of some clear phenotypic patterns (i.e. killer resistance and pseudohyphal growth ability) found in the HS strains tested. However, taking into consideration that this analysis implied a reduced number of strains, we decided to perform a wider genomic survey in order to identify mutations that significantly co-occurred with *IRC7^s^* allele at a population level, discussing the potential role of these mutations in the dominance of HS strains in the wine clade.

### Distribution of IRC7-associated mutations in S. cerevisiae strain populations

Using the genome dataset of Gallone *et al.* (2016), we have studied the distribution of the above-described potential *IRC7*-associated mutations (Table S3) among 150 *S. cerevisiae* strains representing Wine, Sake/Asian, Bread/Mixed, Beer1 and Beer2 populations (Supplementary File S3). Figure 5 shows the global distribution of the previously identified mutations, using *S. cerevisiae* VL3 as the reference (HF) genome. First, we observed that some mutations appeared widespread distributed among all the strains studied (affecting *BBC1, UTP18, PAU6, YML083C,* and *VPS10* genes), so we hypothesised that they were just very specific allelic variants of the reference genome used for the variant calling. Contrary, other mutations were found to be conserved in the Wine clade and mostly lacked in the other clades, appearing as potential hallmarks of domestication in wine populations (affecting to *APC4, VHS1, SDH1, RSF1, MAP1, YDR185D* and *YJL163C* genes) (Figure 5, Supplementary file S3). In addition, we identified mutations with significant co-occurrence rates [across clades (Table S4) and in the Wine clade (Table S5)] with *IRC7^s^* allele.

**Figure 5:**
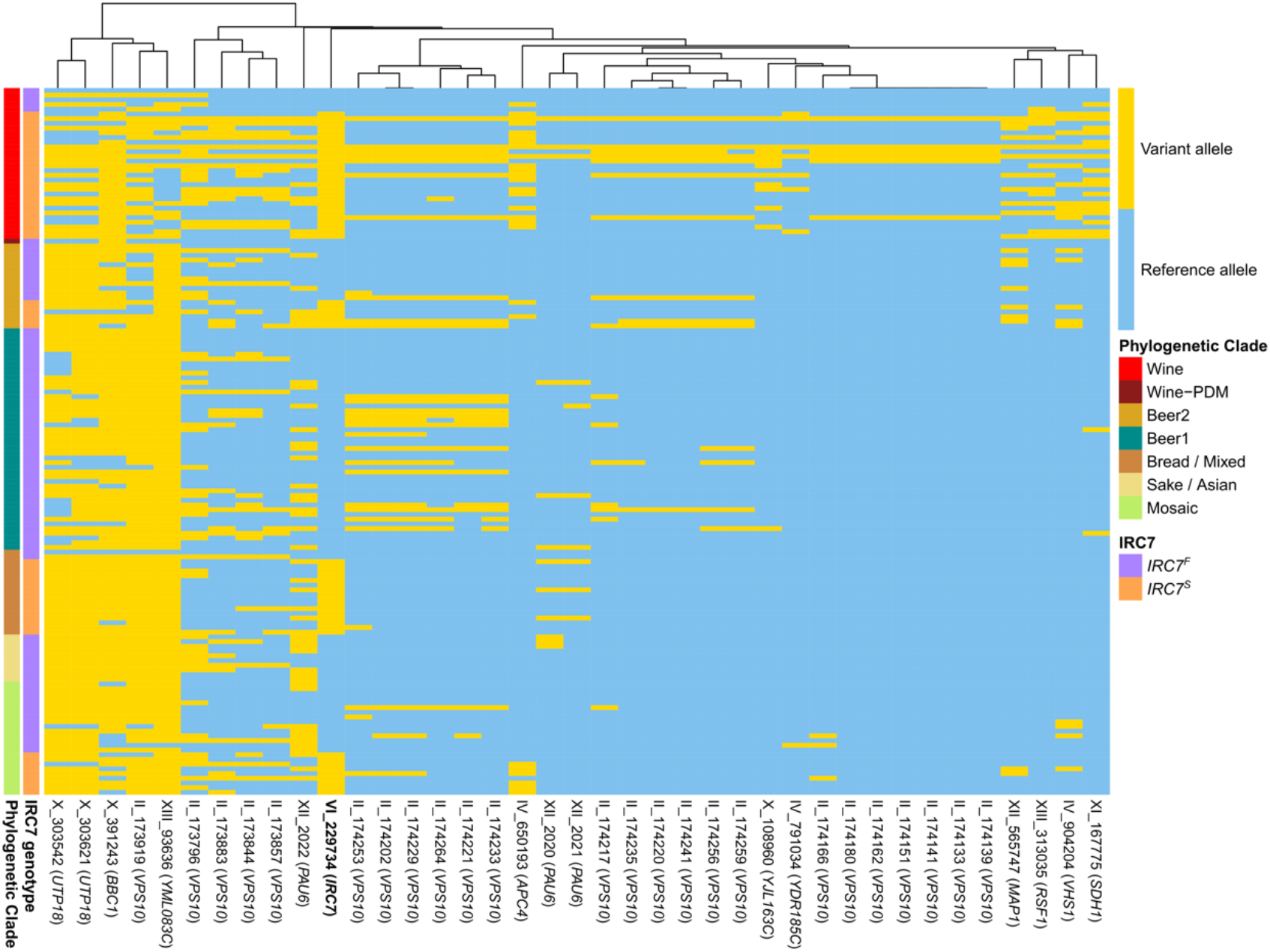
Distribution of the 39 mutations (affecting to 13 different genes) identified in the variant calling analysis and potentially associated to *IRC7* mutation. The presence/absence (variant allele/reference allele) of each mutation in the strain collection (representing six domesticated population) is indicated by the heatmap representation. *IRC7* allele distribution and phylogenetic clades of the strains are highlighted. Strains (represented as rows in the heatmap) are grouped by their phylogenetic clade and *IRC7* genotype. Mutations (represented as columns in the heatmap) are clustered in the dendrogram based on its distribution among the analyzed strains. Supporting data for this heatmap is described in the Supplementary File S3.

Table S4 highlights some mutations with a significant probability of co-occurrence with *IRC7*^S^ allele, independently on the clade, and affecting to the following genes: *APC4, RSF1, YJL163C, SDH1, MAP1, VHS1* and *VPS10. SDH1* and *RSF1* are genes involved in yeast respiratory metabolism, encoding *SDH1* for a flavoprotein subunit of the succinate dehydrogenase, which oxidizes the succinate in the TCA cycle. It has been reported that, during fermentation, TCA cycle is interrupted at the level of SDH complex, but succinate is still formed by the oxidative branch of the TCA pathway (Camarasa *et al.*, 2003). The transition to a respiratory metabolism and the maintenance of certain degree of respiration during fermentation have a direct impact on ethanol consumption at the end of the fermentation, and therefore, on the competitive performance during wine fermentation (Gasmi *et al.*, 2014). Further studies are necessary to understand the potential role of *SDH1* and the mutation we reported here on the metabolism of succinate and the maintenance of respiratory activity during wine fermentation.

Interestingly, Gerke *et al.* (2009) reported that *RSF1* mutation is lacked in wild strains with a high-sporulation efficiency, as oak strains, but occurs in most of strains isolated from vineyards. Our results reinforced this observation, proving that *RSF1* mutation is only found in *S. cerevisiae* Wine clade strains (Figure 5, Supplementary File S3). In *S. cerevisiae* wild strains, sexual reproduction -and therefore sporulation lifestyle-is favored, rather than domesticated strains which present a predominant asexual lifecycle (Liti, 2015). Thus, although further studies are necessary to demonstrate the increased ability of *RSF1-*mutated strains to survive and compete in wine fermentations, our results confirmed this allele variant as a hallmark of domestication in wine yeasts. We should also highlight the mutation detected in *MAP1* gene, due to its functional relationship with Irc7p, as both participate on methionine metabolism. The metabolism of methionine in wine fermentation has a great importance in both fermentation kinetics -increasing the assimilable nitrogen pool during fermentation- and wine flavor -as it is related to H2S off-flavor production (Guitierrez *et al.*, 2013). In the same line, *YJL163C*-putative protein seems to be involved in the iron uptake and transport in the cell.

Finally, some of the above-mentioned mutations are specially found as co-occurring with *IRC7*^S^ strains in the wine clade (Table S5). Apart from the reported role of *VPS10,* the *APC4* gene encodes for a ubiquitin ligase involved in the anaphase inhibitors degradation and the reduction on its functionality could generate an increased competitive fitness (Breslow *et al.*, 2008). *VHS1* encodes for a protein kinase activated by glucose availability, and it has been described as a member of the fermentome group (genes required to accomplish wine fermentation in *S. cerevisiae*) (Walker *et al.*, 2014). *BBC1* gene is involved in the assembly of actin patches, and its deletion results in an improved fermentation performance (Peter, 2017)

The individual effect of all these genetic variants could explain some of the phenotypic observations of this work. However, as it can be inferred from the high co-occurrence frequencies found in *IRC7*-HS-strains, we hypothesize that these genetic variants jointly define a genetic background that could explain the surprising phenotypic characteristics of these strains.

Although further studies will be necessary to understand the phenotypic implications of these identified mutations, we highlight the contribution of the reported findings in the explanation of the outstanding distribution of the *IRC7* alleles. *IRC7* has been deeply studied because it is one of the most important genes related with the production of aromatic thiols that characterize some white wines varieties. However, its paradoxical distribution among the *S. cerevisiae* strains was barely understood until now, and some unproved hypothesis was just proposed. In this work, the extensive phenotypic and genomic conducted studies allowed us to understand the global distribution of the *IRC7* allele among the *S. cerevisiae* linages, to know the associated fitness of the strains according to their *IRC7* genotype, and to identify the *IRC7^S^* accompanying mutations that could explain the dominance and fitness advantage of the strains harboring this *IRC7* allele in wine environment.

## Experimental Procedures

### Yeast strains

*Saccharomyces cerevisiae* strains used in this study (Table S1) were from CYC (Complutense Yeast Collection, Madrid, Spain) and Agrovin S.A. (Alcázar de San Juan, Spain). Sabouraud medium (Oxoid, Hampshire, UK) was routinely used for handling of the strains.

### IRC7 genotyping

Previously, our research group performed the *IRC7* genotyping by PCR analysis (Roncoroni *et al.*, 2011) of a vast collection of *S. cerevisiae* wine strains (Belda *et al.*, 2016). Continuing this work, we performed an extensive *IRC7* genotyping using the genome information of 283 *S. cerevisiae* strains, representing different origins and phylogenetic clades. A local BLAST database was set up for each genome and, using BLASTN searches (1e-4 *E-value* cut-off), we performed the *IRC7* genotyping of the strains. Using *IRC7^F^* sequence of VL3 *S. cerevisiae* strain (available at *Saccharomyces Genome Database*) as a query, it was possible to distinguish the *IRC7* allele that each strain harbors. Strains harboring *IRC7^F^* allele showed ≥99,9% identity, meanwhile stains harboring *IRC7^S^* allele presented 98,5% of identity. This identity percentage difference corresponds with the 38-bp deletion fragment. Strains used in this genotyping analysis, classified in regard to its *IRC7* genotype, are listed in Supplementary File S1.

### IRC7-phenotyping study

#### High throughput phenotyping

We developed a phenotyping screening to characterize the fitness of the strains, in order to find growth ability patterns associated to *IRC7* genotype. Thirty *S. cerevisiae* strains (ten strains belonging to each *IRC7-genotype* group as described lately) were subjected to a high throughput phenotyping study (Table S1). Strains were precultured during 48h in 300 μL of Synthetic Defined (SD) medium (Warringer *et al.*, 2011) with some modifications (2% glucose, 0.14% Yeast Nitrogen Base (BD Difco™, U.S.A.), 0.5% ammonium sulfate, 2,27% succinic acid disodium salt and 0.077% Complete Supplement Mixture (CSM, MP Biomedicals™, U.K.); pH was set to 5.8). After that, strains were inoculated, at a final OD_600 nm_ of 0.2 into 300 μL of the specific SD medium. Assays were performed by triplicate in 96-well plates. All media used are listed in Table S2. The carbon source class indicates that 2% of glucose was substituted with the indicated concentration of the specified carbon source. Likewise, nitrogen source class indicates that 0.5% of ammonia sulfate was substituted with the indicated concentration of the specified nitrogen source. Strains were cultured during 60h and OD_600 nm_ was measured every 4h using a microplate reader (Varioskan Flash Multimode Reader, Thermo Scientific, U.S.A.). Raw OD data were processed as following: OD measure of noninoculated media were subtracted and then, non-linearity of high-density cultures were corrected using the formula OD_correc_ = 0.2453(OD_obs_)^3^ + 0.2735(OD_obs_)^2^ + 0.9779(OD_obs_) – 0.0577 (Warringer & Blomberg, 2003). Finally, wrong data were removed to obtain smoothed growth data, making easy the growth parameters to be extracted. Fitness of the strains was analyzed by extracting the growth parameters (lag time, growth rate and efficiency) from the growth curves using GrowthRates R package (Hall *et al.*, 2014). Growth curves were adjusted to a Baranyi model (Baranyi & Roberts, 1994). Growth parameters, extracted from growth curves, for each strain and condition are shown in Supplementary File S2. Lag phase determination was no possible in all the growth curves (red data in Supplementary File S2), because of the curve shape showed in some media. These missing data were not included in the statistical analysis, neither in the heat map representation.

#### Microvinifications assays

The thirty wine strains were assayed in microvinification assays to characterize the effect of *IRC7-genotype* on wine parameters. Firstly, strains were precultured during 48h in YNB-G medium (0.17% Yeast Nitrogen Base (BD Difco™, USA) and 2% glucose) in 15 mL flasks with 10 mL of medium at 28°C with shaking (120 rpm). Then, strains were inoculated at a final cell concentration of 10^6^ cells/mL in Synthetic Grape Must medium described by Henschke & Jiranek, 1993, that mimics a natural grape must, supplemented with 300 mg/L of sodium metabisulfite and adjusted to pH 3.5. Assays were performed by triplicate in 30 mL flasks with 25 mL of must. Fermentations were performed at 25 °C with shaking at 120 rpm. Once fermentations finished, cultures were centrifugated at 7 000 rpm for 10 minutes to remove biomass. Then, supernatants were stored at −20°C until further analysis. To quantify basic parameters of finished fermentation we used the near infrared spectroscopy method, utilizing one monochromator instrument, Foss NIRSystems 6500 SY-I (Silver Spring, MD, USA). Ethanol content, residual sugars, density, pH, malic acid, lactic acid, volatile acidity and total acidity were determined by this method.

#### Competition assays

To confirm the effect of *IRC7-genotype* on the fitness of the *S. cerevisiae* strains, competitions assays were carried out where two strains, harboring the two different version of *IRC7,* were inoculated in mixed cultures. Four HF-strains (HF1, HF2, HF8 and HF9) and three HS-strains (HS4, HS6 and HS9) were selected for the pairwise competition assays. These experiments were performed into three different media to simulate wine fermentation, beer fermentation and non-fermentative condition. For that, Synthetic Grape Must (Henschke & Jiranek, 1993) supplemented with 300 mg/L of sodium metabisulfite, Beer Must (malt extract 13.2 %, pH 5.2) and non-fermentative medium (glycerol 6%; YNB without amino acid and ammonium sulfate 0.017% (BD Difco™, U.S.); CSM 0.077% (MP Biomedicals™, UK); ammonium sulfate 0.5%) were used as culture media. Cultures were performed, by triplicate, in 40 mL flasks with 30 mL of the correspondent medium, and incubated at 25 °C under orbital shaking at 120 rpm. Previously, strains were precultivated during 48h in YNB-G medium. After that, strains were inoculated in the proportion 1:1, reaching a final concentration of 2 x 10^6^ cells/mL. Weight loss were monitored to determine the end of the fermentation. Final time samples were taken, serially diluted and plated to obtain colonies after incubation at 28°C. Ten colonies were selected from each replicate, and the implantation of the strains was monitored by DNA extraction and PCR by the interdelta polymorphism fingerprinting method (Legras & Karst 2003). Thus, the implantation percentage of each strain at the end of the culture were calculated.

#### Killer activity assays

Killer activity and killer sensitivity of the thirty strains of the study (representing the three *IRC7-genotype* group) were measured using the method described by Santos *et al.*, (2009). Strains to be tested for killer activity were inoculated in ~1 cm diameter concentrated zones onto YMA-MB plates (1 % glucose, 0.3 % yeast extract, 0.3 % malt extract and 0.5 % proteose peptone, supplemented with 30 mg/L of methylene blue, 3 % NaCl and 2 % agar) previously seeded with a lawn (5.0×10^5^ cells/ml) of the strains to be tested for killer sensitivity. Plates were incubated for a week at 20 °C. After that, killer activity was detected by the observation of the halo of inhibition.

#### Oxidative stress evaluation

To compare the oxidative stress level in fermentation conditions between both *IRC7*^F^ and *IRC7*^S^ genotype group, intracellular ROS levels were evaluated. As described in the high throughput phenotyping, strains were precultivated and inoculated in SGM medium. After 24 h of fermentation, cells were treated during 90 min with menadione 1 mM. Then, cells were treated with dihydrorhodamine 123 (DHR 123, Sigma-Aldrich) at a final concentration of 7.5 μg/mL, and incubated in the dark during 90 min at 28°C under orbital shaking. After that, cells were pelleted, washed and resuspended in PBS. Then, the oxidative stress was analyzed by quantified green florescence emission (540 nm) after excitation (485 nm) in a microplate reader Varioskan Flash Multimode Reader (Thermo Scientific).

#### Pseudohyphal growth test

For pseudohyphal growth development, yeasts were grown on minimal medium (0.17% yeast nitrogen base without amino acids and ammonium sulfate, 2% glucose, and 10 mM ammonium sulfate) for 16 hours at 28 °C. After that, cells were harvested and diluted (10^6^ factor). From these dilutions, 100 μL were taken and spread onto solid Synthetic Low-Ammonium-Dextrose (SLAD) medium (0.17% yeast nitrogen base without amino acids and ammonium sulfate, 2% glucose, and 50 μM ammonium sulfate). Plates were incubated for 5 days at 28 °C and colonies were observed by microscopy (10x).

### Genome sequencing and identification of IRC7-associated mutations

Nine *S. cerevisiae* wine strains genomes, in representation of the three *IRC7* genotypes, were sequenced (GenBank accession PRJNA646611). Total genomic DNA was extracted using ZR Fungal/Bacterial DNA MiniPrep Kit (Zymo Research®, USA). Library preparation was performed by enzymatic fragmentation using Nextera DNA Library Prep kit (Illumina, USA). Libraries were sequenced Illumina NextSeq 500/550 v2.5 per Kits. The sequencing coverage was 50X.

Sequence was filter with Trimmomatic v.0.38 (Bolger *et al.*, 2014) with the following parameters: sliding window 15, minimum quality Q20 and minimum length 140. Filtered sequence was aligned with BWA (0.7.15-r1140) (Li & Durbin, 2010) against *S. cerevisiae* VL3 genome (GenBank accession GCA_000190235.1_ScVL3_v01). Variant calling process was performed using GATK4 (4.0.4.0) (McKenna *et al.*, 2010). The process includes: indel realignment, duplicate removal, and performed SNP and INDEL discovery. The parameter was set according to GATK Best Practices recommendations. SNP and INDEL functional annotations were performed by SnpEFF4.3t (Cingolani *et al.*, 2012).

Variant calling results were treated with R version 3.6.3 (R Development Core Team, 2019) and the packages vcfR (Knaus *et al.*, 2017) and tidyverse (Wickham *et al.*, 2019). Bipartite network was built selecting the combination of chromosome, position and alternative and linking with the strain. The resulting network was visualized using Gephi software version 0.9.2. (Bastian *et al.*, 2009).

### Genomic survey

In order to study the distribution of the previous identified mutations across the *S. cerevisiae* population, genomic results were combined with all genomes from the publication of Gallone *et al.*, (2016) (including *S. cerevisiae* genomes from six different phylogenetic populations). In order to compare and combine the results original fastq sequences were downloaded from ENA database. All samples were processed with the same workflow as described above (except for the functional annotation). All tables were processed using R and tidyverse packages. Mutation heatmap was performed using pheatmap (Kolde, 2019).

The co-occurrences with *IRC7^S^* allelic variant of each mutation were obtained using the results from mutations distribution data (Supplementary File S3). The probability of each co-occurrence was calculated following the probabilistic method described by Veech et al. (2013).

### Statistical Analysis

Statistical analysis was performed with the package stats of R software, version 3.6.3 (R Development Core Team, 2019). Analysis of variance (ANOVA) and Tukey post-hoc tests were applied to compare means of the different assays. Principal Component Analysis (PCA) were applied to analysed wine basic parameters of microvinifications assays.

## Acknowledgments

Funding for the research in this paper was provided by Agrovin S.A, under the framework of the CDTI-financed project IDI-20160102, by Pago de Carraovejas S.L.U. CDTI-financed project IDI-20160750 (Centre for Industrial Technological Development, Spanish Ministry of Economy, Industry and Competitiveness, Spain), by the Madri+d foundation and the Rey Juan Carlos University (R&D projects for young researchers under the competitive project ‘WiNetworks’) and by the Ministry of Science and Innovation of Spain (MICINN project: PID2019-105834GA-I00).

Javier Ruiz acknowledges to the Complutense University of Madrid for his doctoral grant (CT17/17 - CT18/17) and Miguel de Celis acknowledges to the Spanish Ministry of Economy, Industry and Competitiveness for his doctoral grant (BES-2017-080024) in the framework of the project CTM2016-76491-P.

