## Supplementary Material for "New insights into the paradoxical distribution of *IRC7* in *Saccharomyces cerevisiae* and its associated phenotypic and genomic landscapes"

Javier Ruiz et al.

### Supplementary material:

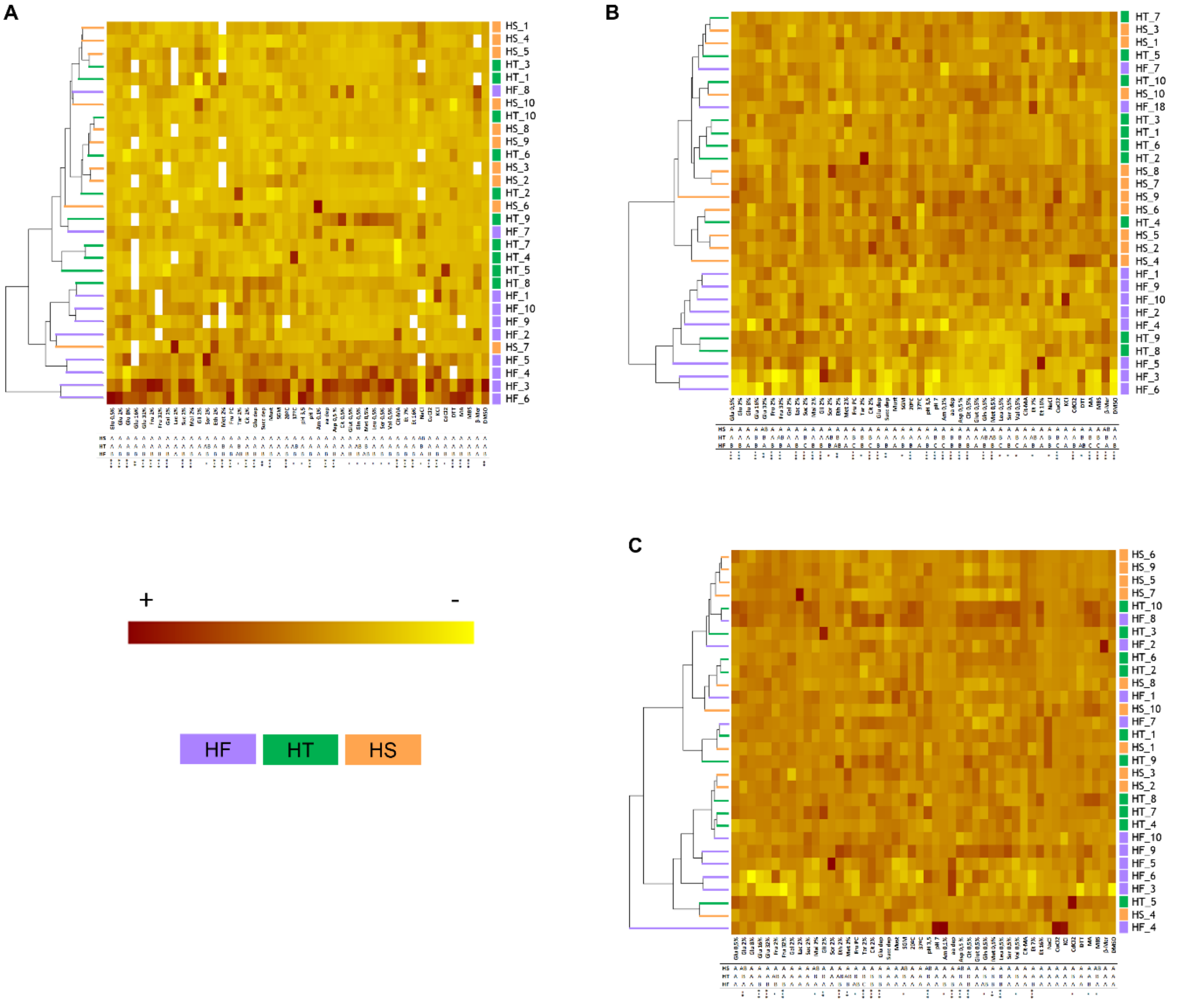


**Figure S1:** Growth ability of thirty *S. cerevisiae* strains, representing each *IRC7* genotype group: homozygous for the full-length allele strains (HF), heterozygous strains (HT) and homozygous for the short-length allele strains (HS). Growth ability were measured by extracting the three growth parameter (**A**: lag time, **B**: growth rate and **C**: efficiency) from the growth curves in each medium of the panel. Strains phenotypic behavior is hierarchically clustered based on growth parameter in the media panel. Growth parameter of the three *IRC7* group of strains in each individual medium were statistically analyzed by an ANOVA and Tukey post-hoc test to compare means. Different letters indicate the existence of statistical differences (* $p < 0.05$; ** $p < 0.01$; *** $p < 0.005$). Missing data are represented by white squares.

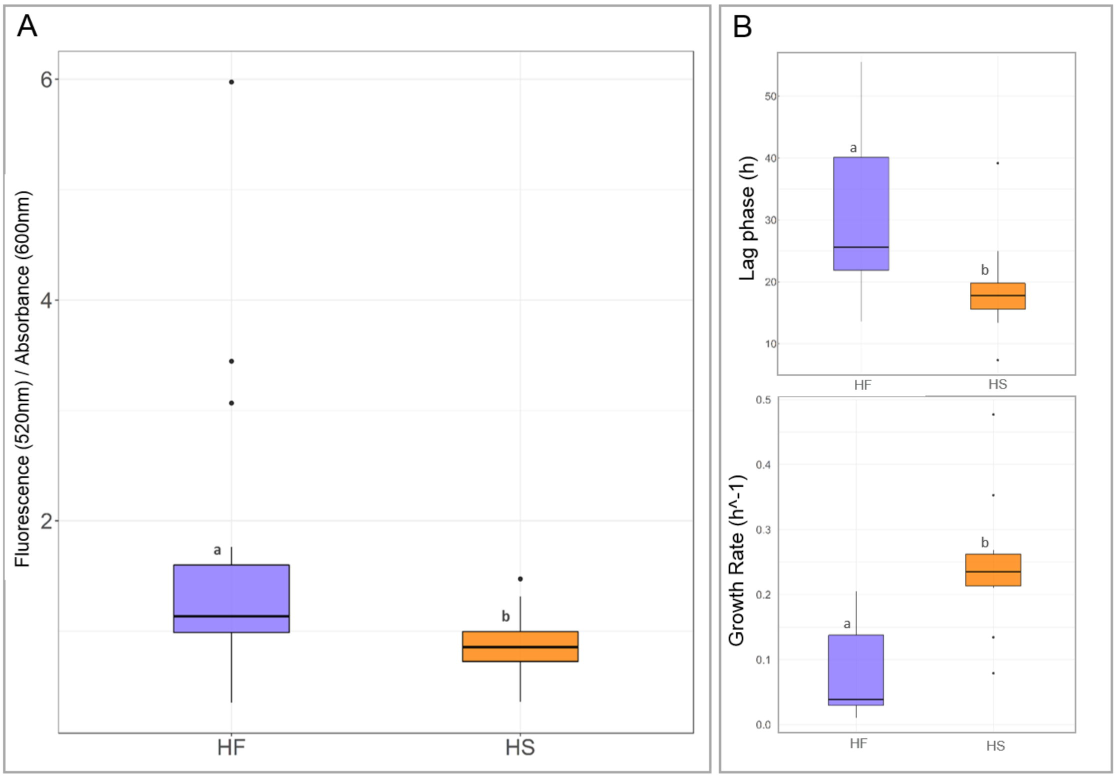


**Figure S2: A** ROS level production were evaluated in the strains after a menadione 2 mM treatment using dihydrorhodamine 123 as indicator. Fluorescence at 520nm were quantified and normalized with the absorbance value of the cultures. Boxplots represent the average oxidative stress level of all the strains of each *IRC7* genotype group. **B** Growth parameters (lag phase and growth rate) were evaluated under the presence of $CuCl_2$ 375 µM. Boxplots represent the average according to their *IRC7* genotype. The performance in all the media panel were consider together in each plot. Different letters indicate the existence of statistical differences ($p < 0.05$).

**A**

| Name | *IRC7* genotype | Pseudohyphal growth |
|---|---|---|
| HS_1 | *IRC7* $^{SL}$ / *IRC7* $^{SL}$ | + |
| HS_2 | *IRC7* $^{SL}$ / *IRC7* $^{SL}$ | + |
| HS_3 | *IRC7* $^{SL}$ / *IRC7* $^{SL}$ | + |
| HS_4 | *IRC7* $^{SL}$ / *IRC7* $^{SL}$ | + |
| HS_5 | *IRC7* $^{SL}$ / *IRC7* $^{SL}$ | nd |
| HS_6 | *IRC7* $^{SL}$ / *IRC7* $^{SL}$ | + |
| HS_7 | *IRC7* $^{SL}$ / *IRC7* $^{SL}$ | + |
| HS_8 | *IRC7* $^{SL}$ / *IRC7* $^{SL}$ | + |
| HS_9 | *IRC7* $^{SL}$ / *IRC7* $^{SL}$ | + |
| HS_10 | *IRC7* $^{SL}$ / *IRC7* $^{SL}$ | + |
| HT_1 | *IRC7* $^{SL}$ / *IRC7* $^{FL}$ | - |
| HT_2 | *IRC7* $^{SL}$ / *IRC7* $^{FL}$ | - |
| HT_3 | *IRC7* $^{SL}$ / *IRC7* $^{FL}$ | nd |
| HT_4 | *IRC7* $^{SL}$ / *IRC7* $^{FL}$ | + |
| HT_5 | *IRC7* $^{SL}$ / *IRC7* $^{FL}$ | - |
| HT_6 | *IRC7* $^{SL}$ / *IRC7* $^{FL}$ | + |
| HT_7 | *IRC7* $^{SL}$ / *IRC7* $^{FL}$ | + |
| HT_8 | *IRC7* $^{SL}$ / *IRC7* $^{FL}$ | - |
| HT_9 | *IRC7* $^{SL}$ / *IRC7* $^{FL}$ | + |
| HT_10 | *IRC7* $^{SL}$ / *IRC7* $^{FL}$ | + |
| HF_1 | *IRC7* $^{FL}$ / *IRC7* $^{FL}$ | - |
| HF_2 | *IRC7* $^{FL}$ / *IRC7* $^{FL}$ | - |
| HF_3 | *IRC7* $^{FL}$ / *IRC7* $^{FL}$ | - |
| HF_4 | *IRC7* $^{FL}$ / *IRC7* $^{FL}$ | - |
| HF_5 | *IRC7* $^{FL}$ / *IRC7* $^{FL}$ | + |
| HF_6 | *IRC7* $^{FL}$ / *IRC7* $^{FL}$ | - |
| HF_7 | *IRC7* $^{FL}$ / *IRC7* $^{FL}$ | - |
| HF_8 | *IRC7* $^{FL}$ / *IRC7* $^{FL}$ | + |
| HF_9 | *IRC7* $^{FL}$ / *IRC7* $^{FL}$ | + |
| HF_10 | *IRC7* $^{FL}$ / *IRC7* $^{FL}$ | - |

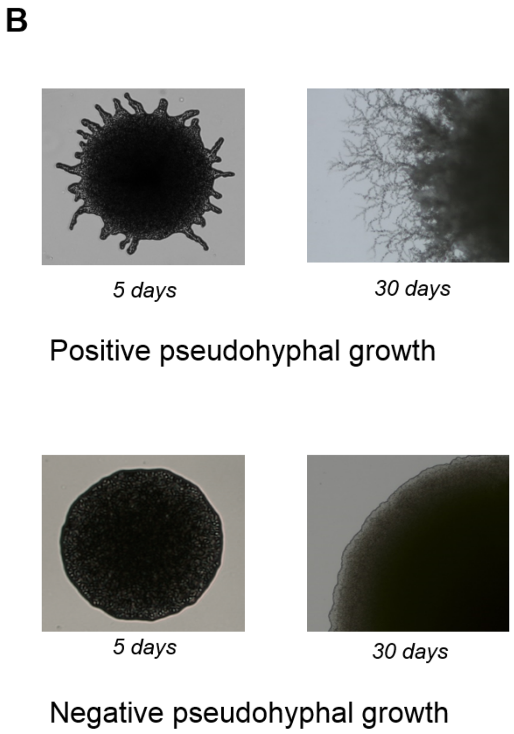


**Figure S3: A** Pseudohyphal growth assay performed in the strain panel (representing each *IRC7* phenotype group) using SLAD medium. nd: not determined. **B** Example results of positive and negative assay (10X microscopy photography).

Table S1: *S. cerevisiae* strains used in the study.

| **Strain** | ***IRC7* genotype** | ***IRC7* group** | **Strain code** | **Origin** |
|---|---|---|---|---|
| S-EM-25 | $IRC7^{SL}/IRC7^{SL}$ | HS | HS_1 | Complutense Yeast Collection |
| S-EM-73 | $IRC7^{SL}/IRC7^{SL}$ | HS | HS_2 | Complutense Yeast Collection |
| S-EM-107 | $IRC7^{SL}/IRC7^{SL}$ | HS | HS_3 | Complutense Yeast Collection |
| S-EM-251 | $IRC7^{SL}/IRC7^{SL}$ | HS | HS_4 | Complutense Yeast Collection |
| S-EM-271 | $IRC7^{SL}/IRC7^{SL}$ | HS | HS_5 | Complutense Yeast Collection |
| S-O-320 | $IRC7^{SL}/IRC7^{SL}$ | HS | HS_6 | Complutense Yeast Collection |
| AG001 | $IRC7^{SL}/IRC7^{SL}$ | HS | HS_7 | Agrovin S.A. |
| AG006 | $IRC7^{SL}/IRC7^{SL}$ | HS | HS_8 | Agrovin S.A. |
| AG008 | $IRC7^{SL}/IRC7^{SL}$ | HS | HS_9 | Agrovin S.A. |
| AG012 | $IRC7^{SL}/IRC7^{SL}$ | HS | HS_10 | Agrovin S.A. |
| S-EM-10 | $IRC7^{SL}/IRC7^{FL}$ | HT | HT_1 | Complutense Yeast Collection |
| S-EM-100 | $IRC7^{SL}/IRC7^{FL}$ | HT | HT_2 | Complutense Yeast Collection |
| S-EM-294 | $IRC7^{SL}/IRC7^{FL}$ | HT | HT_3 | Complutense Yeast Collection |
| S-O-10 | $IRC7^{SL}/IRC7^{FL}$ | HT | HT_4 | Complutense Yeast Collection |
| S-O-166 | $IRC7^{SL}/IRC7^{FL}$ | HT | HT_5 | Complutense Yeast Collection |
| S-O-353 | $IRC7^{SL}/IRC7^{FL}$ | HT | HT_6 | Complutense Yeast Collection |
| AG004 | $IRC7^{SL}/IRC7^{FL}$ | HT | HT_7 | Agrovin S.A. |
| AG007 | $IRC7^{SL}/IRC7^{FL}$ | HT | HT_8 | Agrovin S.A. |
| AG013 | $IRC7^{SL}/IRC7^{FL}$ | HT | HT_9 | Agrovin S.A. |
| AG017 | $IRC7^{SL}/IRC7^{FL}$ | HT | HT_10 | Agrovin S.A. |
| S-EM-115 | $IRC7^{FL}/IRC7^{FL}$ | HF | HF_1 | Complutense Yeast Collection |
| S-EM-129 | $IRC7^{FL}/IRC7^{FL}$ | HF | HF_2 | Complutense Yeast Collection |
| S-O-203 | $IRC7^{FL}/IRC7^{FL}$ | HF | HF_3 | Complutense Yeast Collection |
| S-O-213 | $IRC7^{FL}/IRC7^{FL}$ | HF | HF_4 | Complutense Yeast Collection |
| S-O-331 | $IRC7^{FL}/IRC7^{FL}$ | HF | HF_5 | Complutense Yeast Collection |
| S-O-335 | $IRC7^{FL}/IRC7^{FL}$ | HF | HF_6 | Complutense Yeast Collection |
| AG003 | $IRC7^{FL}/IRC7^{FL}$ | HF | HF_7 | Agrovin S.A. |
| AG015 | $IRC7^{FL}/IRC7^{FL}$ | HF | HF_8 | Agrovin S.A. |
| AG023 | $IRC7^{FL}/IRC7^{FL}$ | HF | HF_9 | Agrovin S.A. |
| WY09 | $IRC7^{FL}/IRC7^{FL}$ | HF | HF_10 | Agrovin S.A. |

Table S2: Culture media used in the *S. cerevisiae* strains phenotyping.

| Medium | Code | Class |
|---|---|---|
| Glucose 0,5% | Glu 0,5% | Carbon source |
| Glucose 2% | Glu 2% | Carbon source |
| Glucose 8% | Glu 8% | Carbon source |
| Glucose 16% | Glu 16% | Carbon source |
| Glucose 32% | Glu 32% | Carbon source |
| Fructose 2% | Fru 2% | Carbon source |
| Fructose 32% | Fru 32% | Carbon source |
| Galactose 2% | Gal 2% | Carbon source |
| Lactose 2% | Lac 2% | Carbon source |
| Sucrose 2% | Suc 2% | Carbon source |
| Maltose 2% | Mal 2% | Carbon source |
| Glicerol 2% | Gli 2% | Carbon source |
| Sorbitol 2% | Sor 2% | Carbon source |
| Ethanol 2% | Eth 2% | Carbon source |
| Methanol 2% | Met 2% | Carbon source |
| Fructose Preculture | Fru PC | Carbon source |
| Tartaric acid 2% | Tar 2% | Carbon source |
| Citric acid 2% | Cit 2% | Carbon source |
| Glucose depletion Preculture | Glu dep | Carbon source |
| Succinic acid depletion | Succ dep | Carbon source |
| Natural Must | Must | Environment |
| Synthetic Grape Must[1] | SGM | Environment |
| Low temperature (20ºC) | 20ºC | Environment |
| High temperature (37ºC) | 37ºC | Environment |
| Low pH (3,5) | pH 3,5 | Environment |
| Low pH (7) | pH 7 | Environment |
| Low ammonia (0,1%) | Am 0,1% | Nitrogen source |
| amino acid depletion | aa dep | Nitrogen source |
| Asparagina 0,5% | Asp 0,5 % | Nitrogen source |
| Citruline 0,5% | Cit 0,5% | Nitrogen source |
| Glutamic acid 0,5% | Glut 0,5% | Nitrogen source |
| Glutamine 0,5% | Gln 0,5% | Nitrogen source |
| Methionine 0,5% | Met 0,5% | Nitrogen source |
| Leucine 0,5% | Leu 0,5% | Nitrogen source |
| Serine 0,5% | Ser 0,5% | Nitrogen source |
| Valine 0,5% | Val 0,5% | Nitrogen source |
| Citruline-Methilamine[2] | Cit-MA | Nitrogen source |
| Ethanol 7% | Et 7% | Toxics |
| Ethanol 16% | Et 16% | Toxics |
| NaCl 0,85 M | NaCl | Toxics |
| CuCl2 375 µM | CuCl2 | Toxics |
| KCl 2 M | KCl | Toxics |
| CdCl2 100 µM | CdCl2 | Toxics |

| | | |
|---|---|---|
| Dithiothreitol 1,6 mM | DTT | Toxics |
| Methilamine 100 mM | MA | Toxics |
| Sodium metabisulfite 400 ppm | MBS | Toxics |
| β-mercaptoethanol 15 mM | β-Mer | Toxics |
| Dimethyl sulfoxide 2% | DMSO | Toxics |

[1] Henschke and Jiranek, 1993, [2] Thibon et al., 2008

**Table S3**: Mutations shared among the HS strains (highlighted in red in Figure 4), identified in the variant calling analysis of 9 of our studied strains (including 3 representative strains of each IRC7-genotype (HS4, HS6, HS9; HT3, HT6, HT10; HF1, HF2, HF9) and VL3 as a reference strain.

| Gene (Sistematic name) | Description | Genomic coordinates | Impact | Codon change | Amino acid change | Effect |
|---|---|---|---|---|---|---|
| *VPS10* (YBL017C) | Type I transmembrane sorting receptor for multiple vacuolar hydrolases | chr02:173796 | Moderate | c.2492A>G | p.Asn831Ser | Missense variant |
| | | chr02:173844 | Moderate | c.2444C>T | p.Thr815Ile | Missense variant |
| | | chr02:173857 | Moderate | c.2431T>C | p.Tyr811His | Missense variant |
| | | chr02:173883 | Moderate | c.2405G>A | p.Ser802Asn | Missense variant |
| | | chr02:173919 | Moderate | c.2369A>T | p.Tyr790Phe | Missense variant |
| | | chr02:174133 | Moderate | c.2155A>G | p.Lys719Glu | Missense variant |
| | | chr02:174139 | Moderate | c.2149G>A | p.Val717Ile | Missense variant |
| | | chr02:174141 | Moderate | c.2147G>A | p.Gly716Glu | Missense variant |
| | | chr02:174151 | Moderate | c.2137T>C | p.Ser713Pro | Missense variant |
| | | chr02:174162 | High | c.2125_2126insTGAAC | p.Ser709fs | Frameshift variant |
| | | chr02:174166 | High | c.2117_2121delCAGTC | p.Thr706fs | Frameshift variant |
| | | chr02:174180 | Moderate | c.2108A>C | p.Lys703Thr | Missense variant |
| | | chr02:174202 | Moderate | c.2086G>A | p.Val696Ile | Missense variant |
| | | chr02:174217 | Moderate | c.2071A>G | p.Lys691Glu | Missense variant |
| | | chr02:174220 | High | c.2067_2068insTA | p.Val690fs | Frameshift variant |
| | | chr02:174221 | High | c.2065_2066delCC | p.Pro689fs | Frameshift variant |
| | | chr02:174229 | Moderate | c.2059A>T | p.Thr687Ser | Missense variant |
| | | chr02:174233 | Moderate | c.2055A>T | p.Lys685Asn | Missense variant |
| | | chr02:174235 | Moderate | c.2053A>G | p.Lys685Glu | Missense variant |
| | | chr02:174241 | Moderate | c.2047A>T | p.Thr683Ser | Missense variant |
| | | chr02:174253 | Moderate | c.2035G>A | p.Val679Ile | Missense variant |
| | | chr02:174256 | Moderate | c.2032G>A | p.Asp678Asn | Missense variant |
| | | chr02:174259 | Moderate | c.2026_2028delCTC | p.Leu676del | Conservative inframe deletion |
| | | chr02:174264 | Moderate | c.2023_2024insCCT | p.Val675delinsAlaPhe | Conservative inframe deletion |

| | | | | | | |
|---|---|---|---|---|---|---|
| *APC4* (YDR118W) | Subunit of the Anaphase-Promoting Complex/Cyclosome (APC/C), a ubiquitin-protein ligase required for degradation of anaphase inhibitors | chr04:650193 | Moderate | c.1751A>G | p.Glu584Gly | Missense variant |
| YDR185C-like protein | Mitochondrial protein of unknown function | chr04:791034 | Moderate | c.481G>A | p.Val161Ile | Missense variant |
| *VHS1*(YDR247W) | Cytoplasmic serine/threonine protein kinase | chr04:904204 | Moderate | c.716G>A | p.Ser239Asn | Missense variant |
| *IRC7* (YFR055W) | Beta-lyase involved in the production of thiols | chr06:229734 | High | c.1023_1060delCGACTCCTTGATTACCCCTGTAAATCCTTGTAATAATA | p.Tyr341fs | Frame shift variant |
| YJL163C-like protein | Putative protein of unknown function | chr10:108960 | Moderate | c.1615G>A | p.Gly539Arg | Missense variant |
| *UTP18* (YJL069C) | Small-subunit processome protein involved in pre-18S rRNA maturation | chr10:303542 | Moderate | c.304A>G | p.Thr102Ala | Missense variant |
| | | chr10:303621 | Moderate | c.225A>C | p.Glu75Asp | Missense variant |
| *BBC1*(YJL020C) | Similar to cell surface flocculin Flo11p | chr10:391243 | Moderate | c.2469_2495delACCTCCAGCACCTCCAGCACCTCCAGC | p.Pro824_Ala832del | Disruptive inframe deletion |
| *SDH1*(YKL148C) | Flavoprotein subunit of succinate dehydrogenase | chr11:167775 | Moderate | c.604C>T | p.His202Tyr | Missense variant |
| *PAU6*(YNR076W) | Member of the seripauperin multigene family, encoded mainly in subtelomeric region. It is active during alcoholic fermentation | chr12:2020 | Moderate | c.156G>T | p.Met52Ile | Missense variant |
| | | chr12:2021 | Moderate | c.155T>C | p.Met52Thr | Missense variant |
| | | chr12:2022 | Moderate | c.154A>T | p.Met52Leu | Missense variant |
| *MAP1*(YLR244C) | Methionine aminopeptidase | chr12:565747 | Moderate | c.53C>T | p.Thr18Ile | Missense variant |
| YML083C-like protein | Protein of unknown function. Strong increase in transcript abundance during anaerobic growth compared to aerobic growth | chr13:93636 | Moderate | c.238G>A | p.Asp80Asn | Missense variant |
| *RSF1*(YMR030W) | Transcriptional activator required for respiratory growth. May interact with transcription factors to mediate the transition to respiratory growth | chr13:313035 | Moderate | c.542A>G | p.Asp181Gly | Missense variant |

**Table S4**: Co-occurrence of mutations with *IRC7*$^S$ allelic variant (c.1023_1060delCGACTCCTTGATTACCCCTGTAAATCCTTGTAATAATA) in *S. cerevisiae* strains across all clades (Supplementary File S3). Only mutations with a probability of co-occurrence higher than 0.95 are indicated and a co-occurrence with *IRC7*$^S$ higher than10 are indicated.

| Mutation | Affecting gene (Systematic name) | Total of mutations | Co-occurrence with IRC7$^S$ | Proportion[1] | Rate[2] | Probability |
|---|---|---|---|---|---|---|
| IV_650193 | *APC4* (YDR118W) | 27 | 26 | 0,96 | 0,45 | 1,00 |
| XIII_313035 | *RSF1(YMR030W)* | 15 | 14 | 0,93 | 0,24 | 1,00 |
| X_108960 | *YJL163C* | 12 | 11 | 0,92 | 0,19 | 1,00 |
| XI_167775 | *SDH1 (YKL148C)* | 20 | 17 | 0,85 | 0,29 | 1,00 |
| XII_565747 | *MAP1(YLR244C)* | 20 | 16 | 0,80 | 0,28 | 1,00 |
| IV_904204 | *VHS1(YDR247W)* | 22 | 17 | 0,77 | 0,29 | 1,00 |
| II_173857 | *VSP10 (YBL017C)* | 29 | 18 | 0,62 | 0,31 | 1,00 |
| II_173883 | *VSP10 (YBL017C)* | 39 | 21 | 0,54 | 0,36 | 0,98 |

[1] Number of co-occurrences with *IRC7*$^S$ by the number of the total mutations.

[2] Number of co-occurrences with *IRC7*$^S$ by total number of *IRC7*$^S$ mutations (58).

**Table S5**: Co-occurrence of mutations with *IRC7*$^{S}$ allelic variant (c.1023_1060delCGACTCCTTGATTACCCCTGTAAATCCTTGTAATAATA) in *S. cerevisiae* Wine and Wine-PDM clades strains (Supplementary File S3). Only mutations with a probability of co-occurrence higher than 0.95 are indicated and a co-occurrence with *IRC7*$^{S}$ higher than10 are indicated.

| Mutation | Affecting gene (Systematic name) | Total of mutations | Co-occurrence with IRC7$^{S}$ | Proportion[1] | Rate[2] | Probability |
|---|---|---|---|---|---|---|
| II_173844 | *VSP10 (YBL017C)* | 13 | 13 | 1,00 | 0,48 | 0,97 |
| II_173857 | *VSP10 (YBL017C)* | 13 | 13 | 1,00 | 0,48 | 0,97 |
| II_173883 | *VSP10 (YBL017C)* | 13 | 13 | 1,00 | 0,48 | 0,97 |
| IV_904204 | *VHS1(YDR247W)* | 13 | 13 | 1,00 | 0,48 | 0,97 |
| IV_650193 | *APC4 (YDR118W)* | 19 | 18 | 0,95 | 0,67 | 0,96 |
| X_391243 | *BBC1(YJL020C)* | 28 | 25 | 0,89 | 0,93 | 0,97 |
| II_173844 | *VSP10 (YBL017C)* | 13 | 13 | 1,00 | 0,48 | 0,97 |
| II_173857 | *VSP10 (YBL017C)* | 13 | 13 | 1,00 | 0,48 | 0,97 |

[1] Number of co-occurrences with *IRC7*$^{S}$ by the number of the total mutations.

[2] Number of co-occurrences with *IRC7*$^{S}$ by total number of *IRC7*$^{S}$ mutations (27).
